# Exploring the roles of temperature and competition in Fennoscandic Holocene plant communities using dynamic sedimentary ancient DNA models

**DOI:** 10.64898/2026.01.06.697859

**Authors:** Marieke Beaulieu, Victor Boussange, Inger Greve Alsos, Loïc Pellissier

## Abstract

Ecosystems are constantly responding to shifting environmental conditions, and the observed structure of ecological communities at any given discrete observation timepoint is part of a larger dynamic response. However, we lack long-term time series of ecosystem responses allowing understanding of the processes driving community dynamics. Sedimentary ancient DNA (*sed*aDNA) offers novel opportunities to link empirical data with ecological processes, providing near continuous records of plant and animal community changes over millennia. Here, we analyzed metabarcoding data from 10 lakes from Northern Fennoscandia characterizing plant and mammal communities at temporal resolution of 100 years over several millenia. We used *sed*aDNA data to test how biotic processes, including temperature-dependent plant growth, competition, shading, and herbivory, may have driven the dynamics of Fennoscandic Holocene plant communities since the Last Glacial Maximum. We compared the explanatory power of ordinary differential equation-based models simulating community changes and including these processes. The fit of the models improved with the consideration of temperature dependent densification and biotic interactions, most notably shading. When considering individual taxa, the importance of shading and herbivory was greatest for the dwarf shrubs, while tree and shrub taxa were more often best predicted by models without competition. Our study demonstrates how using *sed*aDNA time series together with process-based inverse modelling can uncover the mechanisms of species response to climate change, offering potential for more realistic predictions.

## Introduction

The Earth’s climate is constantly changing, driving the biosphere’s permanent transient state of dynamic response. Ecosystem responses, in turn, have the potential to influence the climate itself. Since the last glacial maximum, climate has been a key driver of vegetation dynamics, especially in the early Holocene (Mottl et al. 2021, Wang et al. 2021). Temperature, a key component of climate, directly shapes species’ ecological niches by regulating their physiology and biochemical processes (Addo-Bediako et al. 2000, Chown and Gaston 2016). Indirectly, climate change, particularly rising temperatures, can alter ecological communities through shifts in biotic interactions, such as transitions from facilitation to competition (Olsen et al., 2016) or food web rewiring (Bartley et al., 2019). For example, arctic and alpine plant species, which have a broad temperature tolerance, may face exclusion due to competition as warming temperatures enable the encroachment of taller vegetation (Sturm et al. 2001, Liu et al. 2022, Garıa Criado et al. 2025). The timing and extent of these changes is thought to depend on temperature-driven growth rates, species interactions and other demographic factors (Angers-Blondin et al. 2018, Andreu-Hayles et al. 2020, Liu et al. 2021). The role of ecological interactions in these dynamics remains poorly quantified, leaving a critical gap in our knowledge of how biotic factors mediate the effects of climate change on ecosystems.

Species within communities do not live in isolation but coexist, modulated by complex biotic interactions. Species vary in their ability to disperse, establish, and persist (Svenning and Sandel 2013, Eidesen et al. 2013, Alsos et al. 2015), which influences the rate at which ecosystems respond to environmental changes as well as their trajectories (Bektas et al. 2024). Taxa-specific lags are shaped by species physiology, demography, the physical environment, and biotic interactions (Alexander et al. 2018, Alsos et al. 2022, Block et al. 2022). This variability helps explain why static contemporary species ranges often fail to align with realized ranges in a changing environment (Maclean and Early 2023, Rubenstein et al. 2023, Lawlor et al. 2024). In addition to theoretical and experimental ecology (e.g., Goodwin et al. 2025), empirical ecology can offer critical insights into how species interactions might shape the future of terrestrial plant communities (Van der Meersch et al. 2025). Accurate forecasting of ecosystem dynamics requires an understanding of process-dependent changes, which in turn demands species-rich datasets collected over extended temporal scales. Although most empirical datasets lack these combined characteristics, sedimentary ancient DNA (*sed*aDNA), which captures taxonomically rich genetic information from sediment profiles, has emerged as a powerful tool to bridge empirical data and ecological processes at watershed scales over millennia (Alsos et al. 2024, Callahan et al. 2025).

Paleoenvironmental records, and in particular *sed*aDNA, allow to track specific biotic groups over spatiotemporal scales relevant to global change. Resolution of temporal dynamics has historically been achieved by tracking pollen, spores, and other microfossils in sediment and ice cores (Birks 2019, Lang et al. 2023). More recently, molecular DNA protocols, including target-probe analysis, metabarcoding and shotgun sequencing, have been applied to ancient sediment samples, generating *sed*aDNA data. *sed*aDNA has offered unparalleled insights into the changes undergone by entire plant communities over the Holocene (Li et al. 2025, Clarke et al. 2019, Rijal et al. 2025, Garćes-Pastor et al. 2025), through increased taxonomic resolution and much broader taxa capture (i.e. most plant taxa do not leave microfossils). Although dependent on the quality and resolution of genetic repositories and the development of suitable targeted primers when adopting a metabarcoding approach, *sed*aDNA has the potential to identify all life forms, yielding detailed multi-trophic representations of ancient ecosystems (Kjær et al. 2022, Capo et al. 2023). Furthermore, *sed*aDNA collected from lakes provides species resolution at the scale of the watershed, a physical entity whose natural boundaries are often used to delimit ecosystems (Alsos et al. 2018, Giguet-Covex et al. 2023, Ataman et al. 2025). Hence, the data provided by *sed*aDNA consists of an unprecedented source of information to improve our understanding of the drivers of biodiversity dynamics over millenia.

By explicitly formulating dynamic causal links between state variables, process-based models are valuable tools for testing mechanistic hypotheses regarding macroecological and evolutionary processes (Enquist et al. 1998, Sitch et al. 2003, Brown et al. 2004, Seidl et al. 2012, Rangel et al. 2018). Nearly a century ago, Volterra introduced the Lotka-Volterra competition model, simulating taxa growth within a community (Volterra 1928). Lotka-Volterra models form the basis of non-linear models, accounting for resource-limited population growth. Growth, defined as a change in biomass, connects various biological scales, from the cell to whole ecosystems (McMahon and Bonner 1983), and is intrinsically linked to many ecological processes, including competition and plant-herbivore interactions (Paine et al. 2012, Hilty et al. 2021). By comparing the explanatory power of different model formulations embedding candidate hypotheses on competition, shading and herbivory, we can provide evidence for key processes driving community structures, providing insights into the biotic interactions that influence local plant succession and biodiversity (Turchin 2003). Despite their potential, the integration of theoretical process-based ecological models with real-world data has historically been challenging. Inverse modelling, consisting of retrieving model parameters and processes from observation data, requires advanced algorithms, more so when considering non-linear models with many state variables and parameters (Turchin 2003). While novel methods, together with increasing computation power, become more widely accessible, the potential of these models, testing their formal dynamics on empirical ecological systems has yet to be fully realized.

Here, we leverage long-term plant and animal *sed*aDNA data from the Holocene in Northern Fennoscandia to quantitatively test candidate processes that may have influenced species turnover in response to a changing climate. We use alternative process-based models that we calibrate against the data to disentangle the linear and non-linear effects of warming on ecosystem assembly through biotic interactions. We build upon a comprehensive dataset of plant and mammal *sed*aDNA from ten lakes in Northern Fennoscandia (Rijal et al. 2021, Alsos et al. 2022, Alsos et al. submitted). Northern Fennoscandia was almost fully ice-covered during the Last Glacial Maximum (Hughes et al. 2016) and *sed*aDNA offers the opportunity to track plant community build-up and dynamics under a dramatically changing climate. During the Early Holocene in this region, taxonomic richness increased rapidly alongside significant atmospheric warming, continuing to rise through the slight cooling of the Middle Holocene until stabilizing over the past three millennia (Rijal et al. 2021). The post-glacial arrival times of plant taxa varied and were influenced by adaptations to temperature, disturbance, and light (Alsos et al. 2022), whereas the terrestrial mammals arrived considerably later than the majority of plant taxa (Alsos et al. submitted).

We investigate the process of community turnover over millennia in Fennoscandia, using inverse modelling on a subset of plant taxa to parameterise lake-level process-models for competing model structures.The plant community models based on a set of Lotka-Volterra (LV) differential equations consider densification (growth), herbivory susceptibility, intraspecific competition and interspecific competition under temperature and herbivory forcings. Investigating the effect of these variables on system dynamics, we expect that temperature-mediated growth will overall improve model fit, notably for larger taxa, while the effect of competition, in the form of shading by larger plant taxa, will be most important for the smaller taxa (dwarf shrubs, forbs, grasses and rushes). The presence of herbivores is expected to potentially limit the growth of forage food while favouring the expansion of less palatable taxa. This effect should more likely be detected in lakes with higher gradients of herbivore densities. Our approach provides a robust framework for understanding how climate-driven changes shape ecosystems over long timescales.

## Methods

### Sediment core sampling and *sed*aDNA analyses

Sediment cores were retrieved from ten lakes in Northern Fennoscandia (Norway=9, Finland=1) between 2011 and 2017 (detailed in Rijal et al. 2021, Figure 1). Lake surface areas varied between 0.8 and 55.3 ha, with maximum depths between 1.2 to 34.8 m. To increase the likelihood of capturing DNA of biota from the entire watershed in the lake sediments, selected lakes came from small watersheds (between 0.1 and 10.6 km2). Retrieved sediment cores were subsampled in the lab at regular intervals, and sediment core sections were dated using ^14^C isotope ratio analysis. Based on the age-depth models developed (see Rijal et al. 2021, Alsos et al. 2022), sediment core ages varied from 16.9 ka to 2.6 ka (Figure 1B). Plant and animal taxa were identified from the metabarcoding of DNA sequences obtained from these dated sections of sediment (Rijal et al. 2021, Alsos et al. 2022, Alsos et al. submitted). aDNA sequences were generated from the extraction and amplification (mostly eight times) of *sed*aDNA using vascular plant and mammal specific primers, specifically the P6-loop region of the plant chloroplast trnL (UAA) intron and a section of the mammalian mitochondrial 16S locus (for full methods see Rijal et al. 2021, Alsos et al. 2022, Alsos et al. submitted).

**Figure 1:**
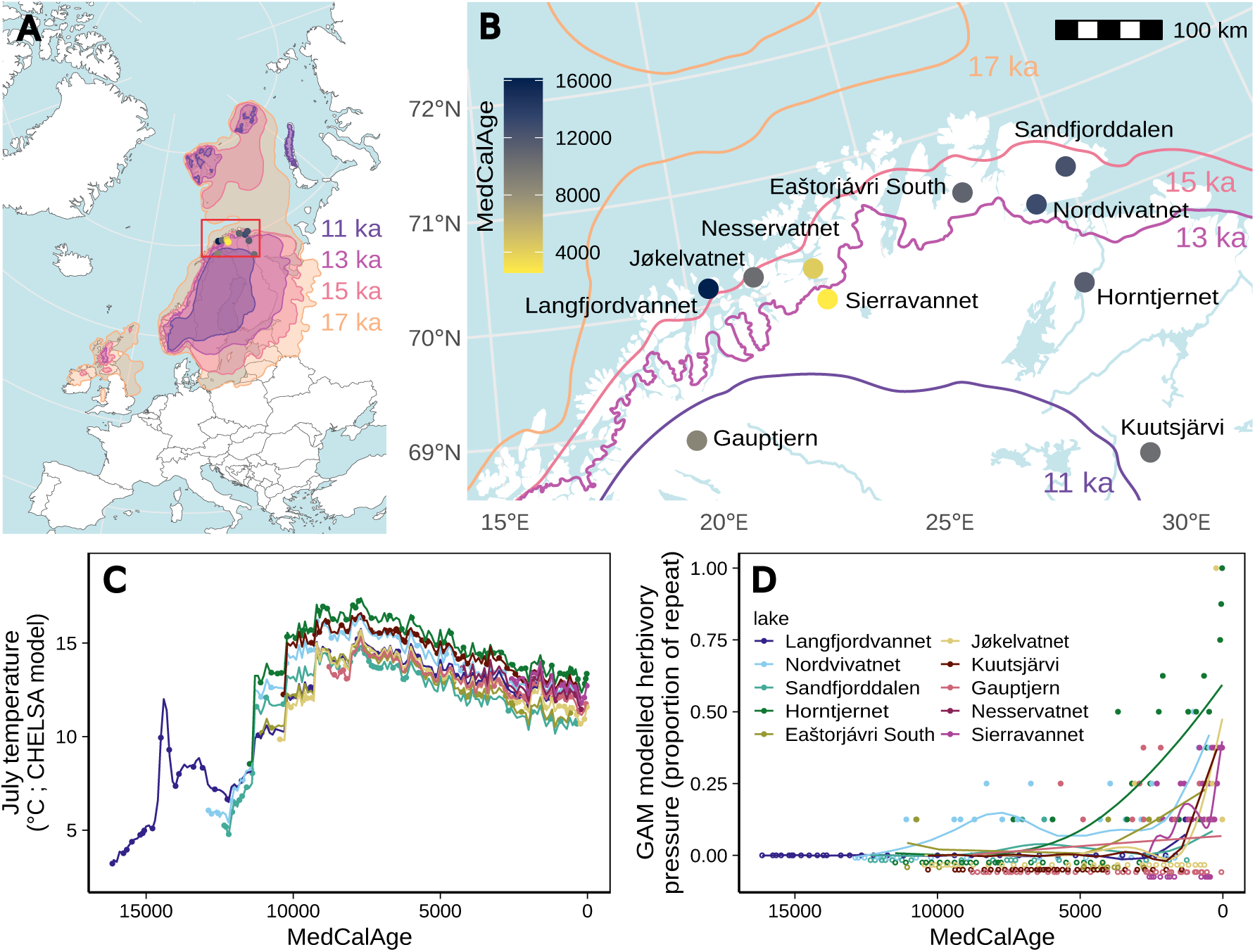
Sampling sites overview. A) Geographic extent of study lake core sites shown in red. Coloured areas represent the estimated extent of the Eurasian ice sheet through time (DATED-1 dataset, Hughes et al. 2016). B) Sampled lakes (as an expansion of study area shown in panel A). Site colours represent sediment core age. Estimated extent of ice sheet margin through time shown as coloured lines. C) Estimated July temperatures for each lake (CHELSA-trace model; Karger et al. 2017, Brun et al. 2022). Points correspond to estimated dating *sed*aDNA samples. D) Reindeer density through time for each lake. Points represent the normalised observed proportion of reads (with zeroes as open points, offset on the y axis). Lines represent the modelled herbivory pressure (GAM).

### Data processing

Data quality and taxonomic resolution can vary between sediment cores due to environmental variability and differences in preservation, most notably for rarer taxa. We first limited the scope of our modelling analyses to plant taxa that were found in 35 samples or more (10% of all samples), retaining 113 of the 315 plant taxa identified. We then focused our initial modelling efforts to 17 taxa chosen for their ecological importance in the European subarctic. These taxa comprised trees and shrubs (*Betula nana/pubescens* [*Betula*], *Pinus sylvestris*, *Saliceae*, *Ribes spicatum*, *Alnus incana*, *Prunus padus*, *Populus tremula*), dwarf shrubs (*Andromeda polifolia*, *Dryas octopetala*, *Empetrum nigrum*, *Vaccinium uliginosum*, *Vaccinium vitis-idaea*, *Vaccinium myrtillus*, *Linnaea borealis*, *Saxifraga oppositifolia* [included among the dwarf shrubs for its somewhat woody branches]), one grass (*Avenella flexuosa*), and one rush (*Oreojuncus trifidus*). We note that although *Saliceae* is treated as a tree/shrub here, taxa within the group are common dwarf shrubs in the region. Although proportion of DNA reads are positively correlated to biomass in the vegetation (Strandberg et al. 2026, Goodall et al. 2025), we here used the more conservative quantification which is number of PCR repeats rescaled from 0 to 1 to avoid any species-specific biases (Alsos et al. 2022).

Mean July temperature estimates at each lake’s coordinates for each sample’s estimated age were interpolated from the CHELSA trace model (Karger et al. 2017, Brun et al. 2022) and used for the temperature forcing in our models (Figure 1C). Temperature data were normalised based on the maximal and minimal temperature values found in the dataset (17.4 °C and -5.3 °C, respectively). We considered herbivory pressure from the perspective of the potential intensity of reindeer (*Rangifer tarandus*) grazing. Repeats data for this taxon were patchier, as is often the case for mammalian taxa, likely due to sparser distributions and mobility of individuals (Murchie et al. 2023). To better capture overall temporal trends, we ran a generalized additive model (GAM) of the normalized repeats over time and utilised the modelled values at each time point (Figure 1D).

### Model structure and fitting

For each lake, we considered eight competing model structures stemming from a base logistic growth model where, for each taxa *i*, the change in the proportion of repeats P depends on its intrinsic densification (growth) rate *g*. Preliminary work demonstrated that models accounting for intraspecific competition, represented by the parameter *c*, which is the taxa’s ability to self-inhibit growth, consistently outperformed all other models. As such, this second parameter is included in our base model (intra [*model 1*]). The carrying capacity for all taxa need not be explicitly considered, as the proportion of repeats data were scaled to values between zero and one. Timesteps were defined by subtracting the oldest date from the regional dataset from all other dates and dividing by 500, with each model timestep covering five centuries. Following the strategy developed by Pellissier et al. (2018), interspecific competition was modelled as the ability of a taxa to tolerate the cumulative biomass of co-occurring taxa, rather than taxa to taxa effects. Intraspecific competition took two forms in our study. In the all-taxa competition model, we considered the cumulative effect of all other species *j* on each taxon *i* which can vary in their susceptibility to competition through the parameter *I* (comp [*model 2*]). In the shading competition model, the parameter *I* was only linked to the cumulative effect of taxa *s* (trees and shrubs, above listed) that can best compete for light (shade [*model 3*]). Finally, susceptibility to herbivory *a* of each taxon *i* was modelled as the response to the modelled proportion of repeats of reindeer *H* (herb [*model 4*]). For these four models we also considered an alternative growth rate structure where growth was represented by two parameters, the afore described *g*, and a second parameter *R*, whose effect on the growth is linearly dependent on temperature forcing variable T (intra temp [*model 5*], comp temp [*model 6*], shade temp [*model 7*], and herb temp [*model 8*]).

Intraspecific competition model (intra)

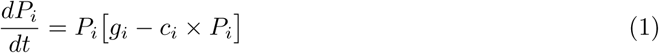

All taxa competition (comp)

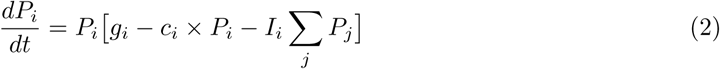

Shading competition (shade)

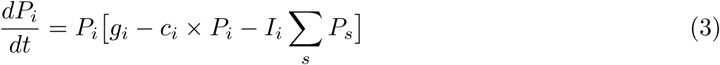

Reindeer herbivory (herb)

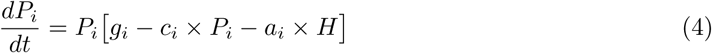

Intraspecific competition, temperature dependent(intra temp)

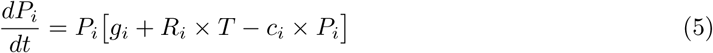

All taxa competition, temperature dependent (comp temp)

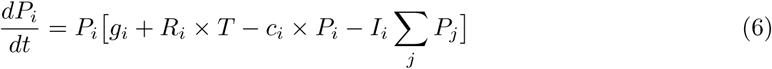

Shading competition, temperature dependent (shade temp)

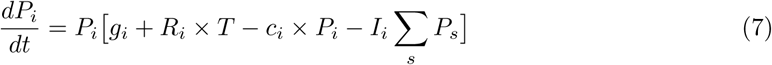

Reindeer herbivory, temperature dependent (herb temp)

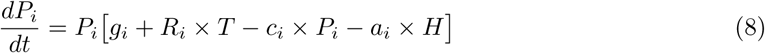

The five most recent data points of each lake were set aside as a validation set. This was a concession, as more sophisticated cross-validation methods would be excessively computationally expensive. For calibration, ordinary differential equation-based models not only require fitting the parameters but also fitting the initial conditions (abundance at the initial time step). Models were specified in Julia using DifferentialEquations.jl (Rackauckas and Nie, 2017). A two-stage method was adopted. First, we used the differential evolution optimiser provided by BlackBoxOptim.jl to find rough estimates of model parameters over 10k iterations, using the L2Loss cost function from DiffEqParamEstim.jI (Rackauckas and Nie, 2017), implementing the sum of squares error (SSE) with first differencing to minimize the weighted cost function (i.e. incorporating the differences of data points at consecutive time points to add more information about the loss function trajectory). In the second step, we minimized the SSE of the models through gradient descent over 10k iterations using the parameters obtained from the derivative-free optimisation as starting parameters. The gradient descent algorithm used was the quasi-Newton optimisation method, BFGS. Quasi-Newton methods, also known as secant methods, are a compromise between inefficient first order gradient descent and computationally costly second order Newtonian methods (Dennis and Moŕe 1977). We limited the scope of our parameter search to values between 0 and 5 for *g*, *R*, *c*, and *I* and between –5 and 5 for *a*, allowing the effect of the presence of reindeer to be positive or negative for a given taxon. Initial parameter values were uniformly drawn within these ranges. As initial conditions may have a strong influence on the fitting process, we computed the initial taxa values as equal to the mean of the first three data points (or 0.001 when the taxa were not detected in that time frame) to account for fluctuations in the observations. For each lake, we iterated through the aforedescribed two-stage method 10 times, as different initial conditions of the parameters may lead to different parameter estimates.

### Model comparison and evaluation

For each lake, a total of eighty models were fitted. To assess method performance and the explanatory power of the fitted competing models, we used a validation-based model selection approach. We first compared the ability of the different models to fit the training dataset. For each lake, we estimated the overall success of the model fitting process of Eq. 1-8 for each iteration by comparing the predictions of the calibrated models to the *sed*aDNA data and calculating the mean square error (MSE).

We then computed the training skill (Skill_Train_)of the model (eq.1) using GAM models (P = s(AgeCalMed, k=5)) fitted on the training dataset as the baseline (Figure 2, S1). Linear models were used when there was insufficient data variation to fit a GAM model. Higher skill values indicate a better fit of the process-based models relative to the GAM models, with positive values indicating improved performance and negative values indicating a poorer fit. We omitted models where the MSE of the baseline model was inferior to 1E-9, as these perfect fits of constant observations create stark outliers in average model Skill_Train_.

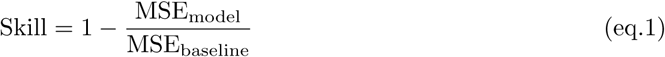

**Figure 2:**
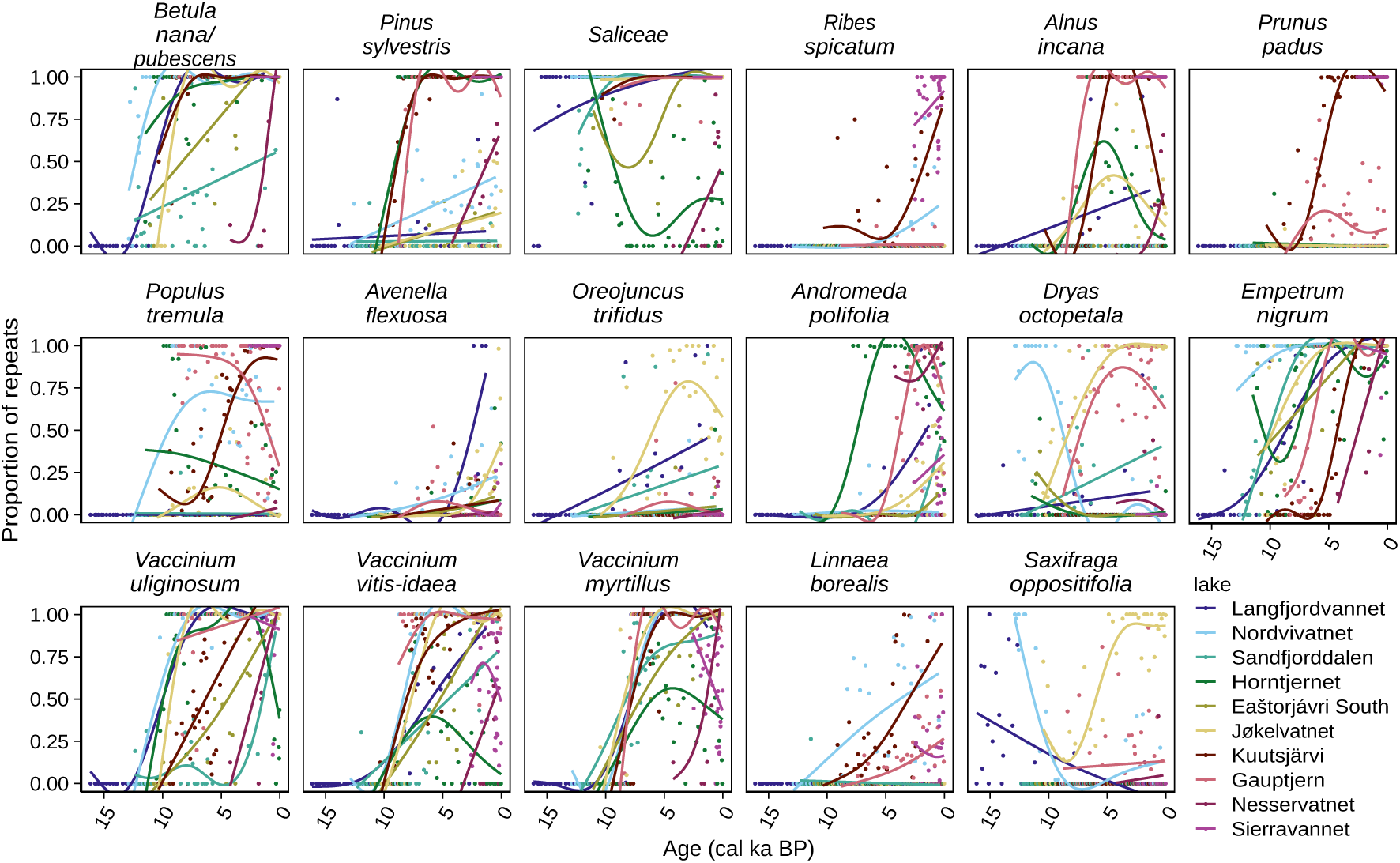
Temporal changes in proportion of repeats for modelled taxa. Points and coloured lines represent respectively the observations and GAM trendline for each lake.

This was done for each taxon and we also averaged over all the taxa to gauge the general performance on a given model run for the entire community. We then considered the top decile of the models based on this fit criterion, distinguishing between models that performed on average better than the baseline model (Skill_Train_¿0) than those that performed on average worse (Skill_Train_ ¡0). For all the top decile models, for both the average taxon response and individual taxa from each lake, we then calculated the residual MSE error on the validation data points for both the process-based model and training dataset GAM model to calculate the Skill of the models in predicting future datapoints (Skill_Test_). This distinguished models that were potentially overfitted and identified potential biases in the estimate of model performance. We nevertheless also report the Skill_Train_ obtained on the training dataset, to evaluate each model’s fit.

## Results

### Temporal abundance trends for individual taxa

Taxa varied in their presence-absence in the lakes and changes in proportion of PCR repeats, although most taxa saw increases in the proportion of repeats at most sites through time (Figure 2). Notable exceptions are *Saliceae* and *Saxifraga oppositifolia*. Several taxa, mainly dwarf shrubs, showed decreases in the proportion of repeats in more recent times, which may be in part linked to decreases in temperature around 7.5 cal ka BP, lake specific increase in pine, and/or increases in reindeer grazing from 5 cal ka BP (Figure 2).

### Lake community model (average taxa response)

Model optimisation success, as estimated by the average taxa Skill_Train_, varied widely across lakes and model optimisation runs (Figure 3A). Across models and model runs, large variances in Skill_Train_ were sometimes observed and this was more pronounced in lakes Kuutsjsavri Gauptjern and Nesservatnet (Figure 3A). While most of the models tested presented similar distributions of residuals, the competition and shade models (with and without temperature dependent growth) showed a larger variance in performance between runs; this was found in most lakes. These results suggest that the performance of our optimisation method is dependent on the values of the initial parameters, and that the strength of this effect depends on the model structure.

**Figure 3:**
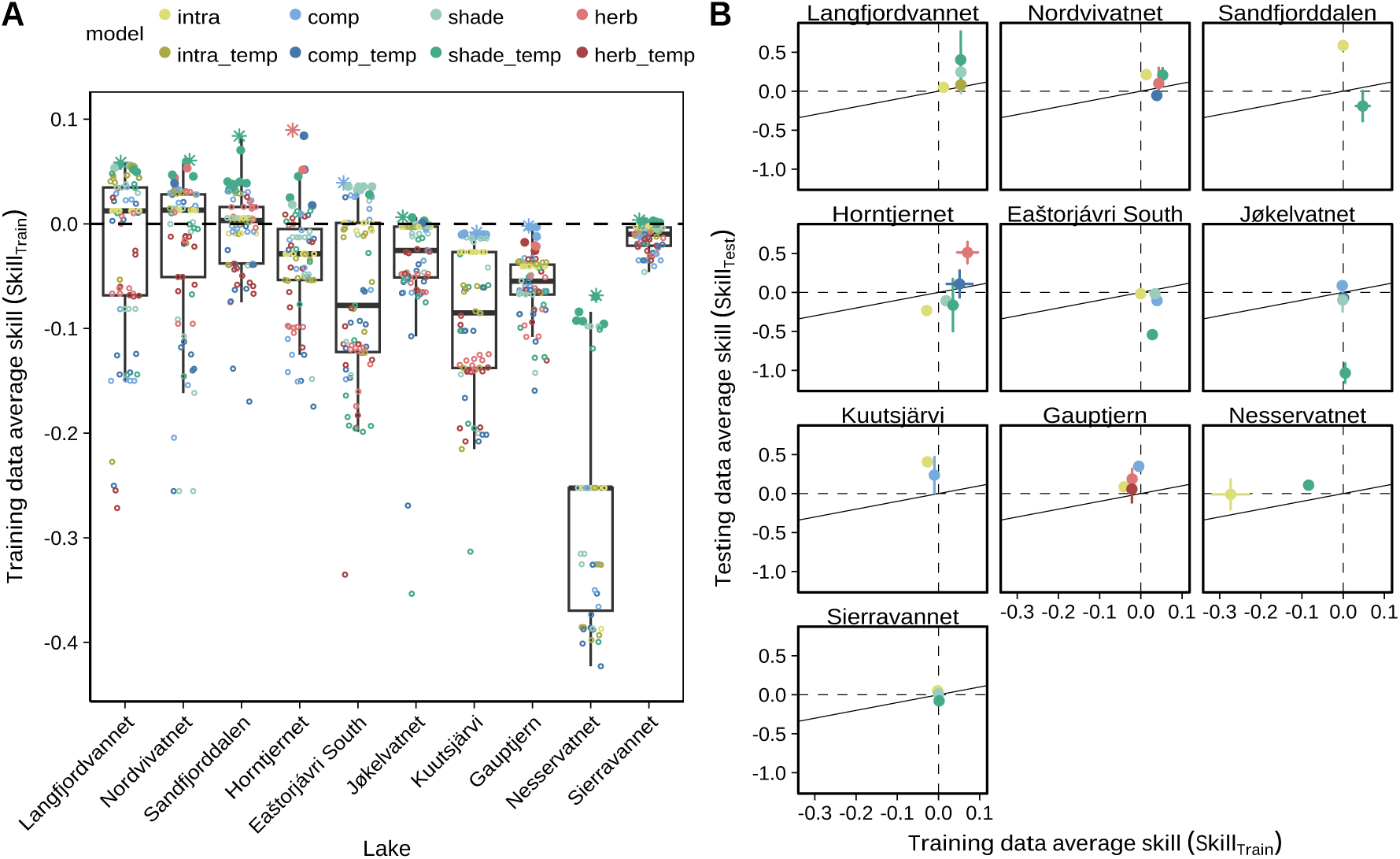
Model fit. A: Community-wide training data fit as represented by the average taxon training skill (Skill_Train_) for each model run (8 competing models over 10 runs, intra = intraspecific competition, comp = All taxa competition, shade= Shading competition, herb= Reindeer herbivory). Large filled points represent the top decile of models for each lake based on Skill_Train_, with the star representing the best fitted model overall. Open circles represent the remaining fitted models. B: Performance of the top decile of average taxa models based on the average Skill_Train_ of the testing dataset for each lake. For each lake, model runs were grouped for each of the model structures, using the same colours as in A). The null model (intra [*model1*]) is represented in yellow for comparison but was only retained in the top decile for five lakes (Eastorjavri South, Jøkelvatnet, Kuutsjsarvi, Nesservatnet, Sierravannet). The full line represents the 1:1 ratio while the dotted lines represent the model skill thresholds for training and testing respectively.

When looking at the models with the best training skill across all taxa, the shade model with temperature-dependent growth performed best in six of the ten lakes, followed by the competition model in three lakes (Jøkelvatnet, Estorjavri South and Gauptjern) and the herbivory model with temperature in one lake (Horntjern, Figure 3A). The latter is also the lake with the greatest gradient in expected herbivory pressure (Figure 1D).

Focusing on the top decile of these lake-level models based on Skill_Train_, we found that, the models that had high Skill_Train_ values also had high Skill_Test_ values and that model performance was adequate (Figure 3B). While, this good performance might also be in part due to the poor predictive ability of the baseline GAM models as the MSE of the test data was consistently greater than that of the training data (S2), we also include the skill of the intra model which allows us to compare each models’ skill with that of the simplest model. We find that, while in most cases the intra model is as good or better at predicting unseen data, interaction models perform better in some lakes. The shade temperature model performed best in Langfjordvannet, the herbivory model in Horntjernet, and the competition model outperforms the alternatives in both Nordvivatnet and Gauptjern. While Horntjernet and Gauptjern had the highest number of taxa with overall decreasing or fluxuating proportion of repeats through time, taxa in Langfjordvannet and Nordvivatnet had simpler and generally increasing trends (Figure 2). This partly supports the idea that increased complexity in taxa dynamics might require complex models. Models in lakes with longer time series generally had better training skills (Figure 3A) but this was not reflected in their testing skills (Figure 3B) and when looking solely at the MSE of the models these models tended to have larger test MSE values (S2). This trend may indicate that changes in the drivers of community dynamics over time are leading to greater mismatches in more contemporary data compared to the older data used to generate the models. Alternatively, inherent structural inaccuracies (process error) which increase when integrated over time (Rosenbaum and Fronhofer, 2023), which account for the poorer predictive ability of models fitted on longer time series. Also, lakes with shorter time frames have shorter intervals and provide more detailed temporal resolution of taxa turnover, which may improve community modelling.

### Taxa-specific models

When considering the performance of the models based on their fitting of the training dataset for individual taxa, constraining for models with (Skill_Train_) values greater than zero, we found differences in the top decile of models best predicting individual taxa, with observable trends when comparing the broader taxonomic groups to which they belong (Trees and Shrubs, Dwarf Shrubs, Grasses and Rushes; Figure 4). Top model groups were unrelated to the general temporal turnover trends of the taxa, with the exception of intra models only being retained for taxa showing overall increasing or constant trends through time. We note however that this absence of pattern might be influenced by the relatively small number of sites compared and the greater occurence of taxa showing overall increasing or generally constant trends through time rather than fluctuating or overall decreasing trends (81% and 19% respectively). Dwarf shrub models were best fitted with the inclusion of susceptibility to shading or herbivory, most often with temperature dependent growth, while trees and shrubs trajectories were best modelled when considering all other taxa, or no community processes at all, with temperature less often considered. Best models for grasses and rushes appeared intermediate to the two other groups (Figure 4). Among the dwarf shrubs, temperature-dependent growth was more often found for the three *Vaccinium* taxa modelled (Figure 4). We note that densification rate was captured by two parameters, one of which was temperature independent, but we found no apparent relationship between these two parameters.

**Figure 4:**
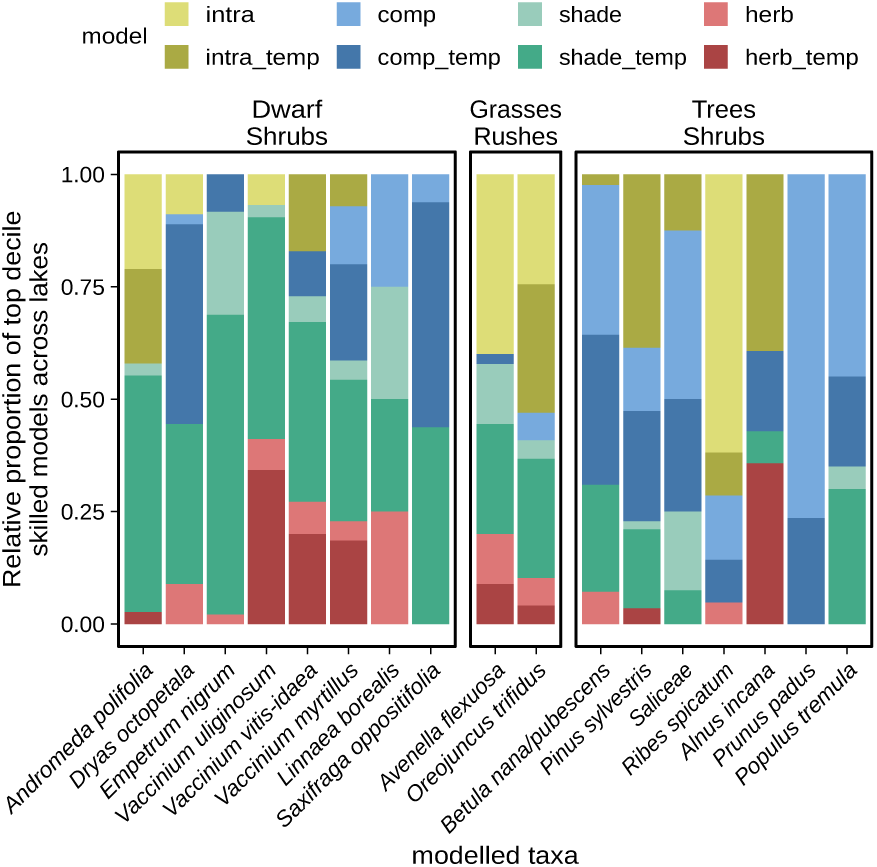
Proportion of each model category retained in the top decile of skilled training models summed across all lakes, omitting models with Skill_Test_ values inferior to 0. Taxa are split into broader groups. Colours represent the four competing interaction models while darker colours represent models that included temperature dependent growth.

The percentage of the top decile of taxa-specific models with Skill_Train_ values greater than zero varied from 14 (*Oreojuncus trifidus*)to 100 (*Alnus incana*), with half of the models retained on average when omitting models with Skill_Test_ inferiour to zero. Overall group trends remained whether these models were omitted or not (not shown). A number of models performed comparatively poorly when tested on unseen data. This was most notable for several of the dwarf shrubs (*Vaccinium uliginosum*, *Vaccinium vitis-idaea*, *Vaccinium myrtillus*; S1, S2). While our best models were generally not overfitted, the taxa-specific null model still outperformed the community interaction models test data in little over half the cases (S3). This could be in part due to the more complex models often not being well constrained to the upper bounds set outside of the training range or because in the last time-steps the effects of parameters important during the transient phase are reduced as the model converges to an equilibrium.

While parameters for competition and shading were limited to positive values (with negative effects on a given taxa’s proportion of repeats data for the latter variables), herbivory pressure was allowed to take both positive and negative values. We found that while herbivory was not often retained as a top decile model for the taxa modelled, when this parameter was retained, it often had a positive effect on tree and shrubs proportion of repeats data, most notably for *Alnus incana* and *Populus tremula* (Figure 5). Although the effect of reindeer was at times more often positive for dwarf shrub species, notably *Vaccinium myrtillus* and *Empetrum nigrum*, generally, the effect of reindeer was negative for dwarf shrubs as well as for grasses and rushes (Figure 5).

**Figure 5:**
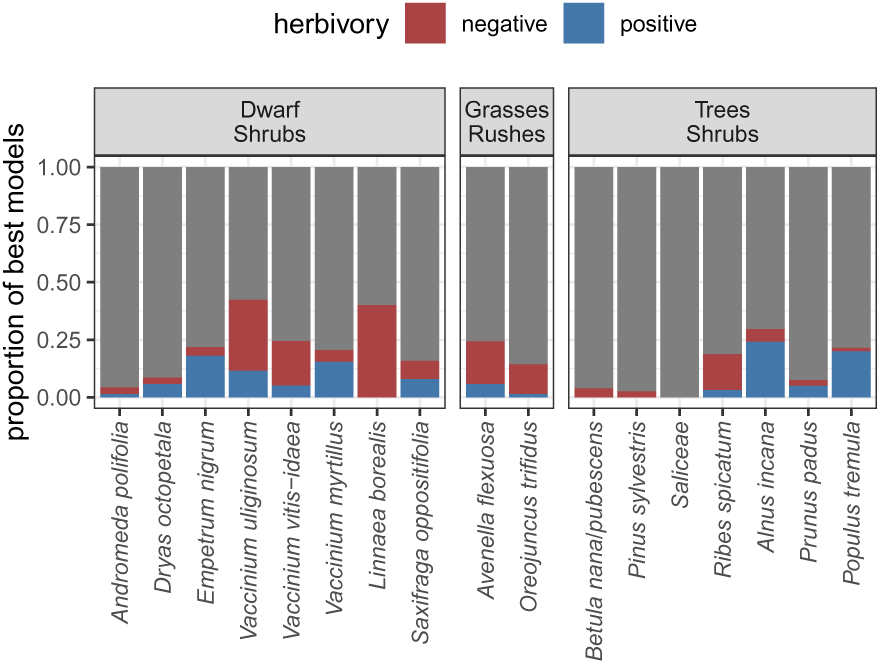
Importance of herbivory in top decile models across lakes grouped by broader plant categories. Colour indicates whether the parameter had a positive or negative effect on the proportion of repeats.

## Discussion

Biotic community responses to temperature changes are a complex interaction between abiotic and biotic drivers (Blois et al. 2013). Process-based models have been previously used to model plant community dynamics over more limited time frames (with yearly to decadal resolution) by incorporating ecological processes, offering insights into drivers of change (reviewed in Decocq et al. 2022). Here, we leverage species records from lake sediment cores, tracking Northern Fennoscandic plant communities over millennia. We found dynamic community processes to consistently explain the data, as null models (intraspecific competition, models 1 and 5) were always outperformed when considering the overall watershed community. Temperature-dependent growth and susceptibility to shading by trees and shrubs best explained the *sed*aDNA proportion of repeats dataset, with competition among all plants or herbivore grazing improving data fitting in some watersheds. Best model fits varied across morphological groups. Shading by larger taxa, temperature-dependent densification and herbivory best explained dwarf shrub temporal dynamics, providing insight into dwarf shrub succession. For trees and shrubs, null models and all-taxa competition more often performed best which suggests that these taxa are less affected by other plant taxa or that our models did not address the community processes relevant to tree and shrub succession. Together, our analyses demonstrate how combining novel *sed*aDNA data with process-based models allow deciphering the mechanisms that drive changes in plant assemblages over time.

Competitive interactions modulate the timing of species response to temperature change (Alexander et al. 2015), and both of these processes were important in the models. We find that among the biotic interactions tested, competition for light, as represented by the response to the density of trees and shrubs, best allowed to describe the succession patterns in the watersheds. This overall trend was principally driven by the sensitivity of dwarf shrubs to the cumulative influence of the larger taxa. Also among the dwarf shrubs, temperature dependent densification was retained. It has been shown that warmer conditions are linked to increased recruitment among dwarf shrubs (Graae et al. 2008). As ongoing warming favours the upwards and poleward expansion of higher canopy taxa (Bjorkman et al. 2018, Garıa Criado et al. 2025), our work suggests that the dwarf shrub communities typical of Northern Fennoscandia may face competition pressuring them from their current range despite warmer temperatures allowing greater densification. Our findings support the hypothesis that novel competitive interactions are likely to affect dwarf shrubs’ responses to climate change (Alexander et al. 2015). In contrast, we find limited evidence that other plant taxa have much of an influence on trees and shrubs, which may be primarily driven by abiotic factors or relationships not addressed in our models. On one hand, it is possible that by considering only the sum of the density of shading taxa or all other taxa we are missing specific taxa-taxa influences. The most notable of these would be the allelopathic effects of *Empetrum nigrum* and other dwarf shrubs (Gonzales et al. 2015, Angers-Blondin et al. 2018). We also note that the scope of the species and interactions considered were limited. For example, tree migration lags have been linked to reduced diversity of ectomycorrhizal fungi partners diversity (Van Nuland et al. 2024) while shifts in the associated microbial community increase the tolerance of trees to climate change (Allsup et al. 2023). Process-models have the potential to find support or disprove important species interactions, such as competition for light, but the choice of relationships to be considered remains daunting.

Temperature is the main environmental parameter that regulates the physiology of an organism (reviewed in Knapp and Huang 2022) and thus should be determinant in community dynamics (Alexander et al. 2015). While temperature was often important, temperature-dependent densification (growth) was not systematically retained in our best models, despite the considerable temperature gradients covered. We found that it was more often among the best models for the dwarf shrubs, which did not support our hypothesis that the larger taxa would be more influenced by temperature. This could be because the tree species were not present in the early cold period, and only arrived after the major temperature increase had taken place. Furthermore, hindcasted climate models may fail to adequately represent local-scale changes, which are likely important to account for in studying range shifts (Lenoir and Svenning 2015). Improving climate models to represent local conditions may only provide limited improvements if they fail to account for climate proximity, that is climate as experienced by the organism itself (Klinges et al. 2024) or even the effect of plant diversity itself on local scale climate (Beugnon et al. 2024). While it remains to be seen how much improvements can be made to hindcasted models, contemporary evidence suggests that improvements to climate models will better address the effects of climate change on community dynamics.

Herbivory and herbivore preference is an important determinant of plant community composition at small scale (Kempel et al. 2015, Koerner et al. 2018), but whether its effect on community dynamics can be detected in lake sediment is unclear. Our models provided some support to some dwarf shrubs, notably *Vaccinium uliginosum* and *Linnaea borealis*, being negatively affected by reindeer densities in the watershed. In contrast, the trees and shrubs *Alnus incana* and *Populus tremula* were more positively affected. Previous review of the long term effects of reindeer on dwarf shrubs have shown these to be weak (Stark et al. 2023) and of little congruence (Bernes et al. 2015). While *Vaccinium uliginosum* is a preferred forage taxa for reindeer (Brathen and Oksanen 2001) and could explain the negative impact of reindeer on its density, we do not observe similar relationships for *Vaccinium myrtillis*, another preferred forage food among the dwarf shrubs. The relatively strong negative impact of reindeer on *Linnaea borealis* was surprising, but can be explained by this plant mainly occurring at Langfjordvannet and Kuutsjsarvi, two sites where reindeer were not found until recent, assumed human-mediated dispersal. Similarly, the positive impact of reindeer on *Alnus incana*, a known preferred forage food for reindeer, could be due to low sample size and reindeer only recently being introduced to Nesservatn, were *Alnus* is found in 8 of 8 replicates of almost every sample. Potentially, there could be a process not included here that correlated with herbivory. While we did not find reindeer herbivory to be important for *Betula* and *Saliceae*, preferred forage among the trees and shrubs, we note that these groups are not yet identifiable to the species level, and more detailed taxonomy might better uncover existing dynamics. Our models might be capturing indirect signals and it would be of great interest to expand the scope of our analysis to include lichens, as this symbiotic group is important in Fennoscandia and is best known to be strongly negatively impacted by reindeer grazing which might help best interpret these results. Furthermore, although reindeer generally represent a greater biomass, the transient effects of reindeer grazing on plant communities could be less important than the local and intense foraging of small mammals (Olofsson et al. 2004).

### Study limitation

While the patterns inferred from the model training are quite clear, their interpretation is tempered when considering the testing data, as, although the performance of the top fitted models was adequate, the null models most often best predicted the test data, both when considering a single model for the entire community or independent models for individual taxa. As it was not feasible to perform true bootstrapping given the number of models to compare and the necessity to test the effect of different initial parameter values, our confidence in our test data results themselves are limited. The test data consisted of the most recent data. Although the limits of the bounded response variable were respected for the training dataset, this did not extend beyond that timeframe, which led to some large divergences, most notably for the herbivory models (S2). Furthermore, it is not currently known if we should expect the community dynamics to be constant through time and/or due to the drivers of community dynamics changing through time (Doncaster et al. 2023). While we cannot exclude that our results are the effect of overfitting, we note that results were quite consistent among the lakes and the taxa, the models still performed well on the test data, and we limited our models to 2-4 parameters to avoid the pitfall of comparing models with larger differences in the number of parameters.

*sed*aDNA data are not without limitations. The amount of DNA released into the environment varies among species, DNA preservation depends on physical substrate characteristics, and biases during DNA amplification (notably during the PCR step) are challenging to correct. Moreover, for common to dominant plant species as several of the dwarf shrubs as well as *Betula* and *Salix* studies here, the conservative quantification using number of PCR repeats may underestimate growth as they often occur in all eight repeats. For these taxa, using proportion of DNA reads may provide a better quantification (Strandberg et al. 2026).

*sed*aDNA data remains noisy and inverse modelling is computationally costly, more so when inputting little prior knowledge about the parameters being retrieved. If we wish to improve the community models to incorporate more realism, increasing sampling frequency from the sediment core could help better resolve temporal trends and better constrain the models. One of the challenges limiting the use of real data to fit parameters of process–based models is the computational cost of calibration methods, which grows significantly as the number of parameters increases. Our current modelling framework does not allow us to incorporate the full diversity of taxa represented in *sed*aDNA datasets. Even when limiting the number of taxa considered (17) and the number of parameters (max 4), each model required the fitting between 17 and 68 parameters. Despite not including any prior expectations in the model fitting, the response variables modelled were proportion of repeats of each taxon which were normalized to values constrained between 0 and 1, this allowed us some confidence in restricting the parameter space explored. Despite this, over 10 runs, models did not consistently converge to values minimizing the model error (Figure 3). This is likely dependent on the combination of the start parameters considered and the model structure. Furthermore, our models did not constrain state variables to proportions in the range [0, 1] beyond the temporal range of the training data. Improving the model’s formulation so that the state variables are better constrained could provide additional inductive bias that will benefit the forecasting ability. We found that it is necessary to repeat the calibration to mitigate for some runs failing to converge to adequate fitting of the date. More advanced calibration methods, such as partitioning the data (Boussange et al. 2025) could improve the robustness of the calibration. Our modelling approach relied on few priors, which is appealing with regards to versatility (e.g. for studying less-known taxa), but the inclusion of known or expected ecological dynamics might improve its performance. Hybrid dynamic modelling, where neural networks are used within process-based models as flexible terms to capture complex dependencies, offer interesting venues to improve explanatory power and forecast skills of community dynamic models.

Biotic interactions impact individual species distributions and are discernible at macroecological scales (Araújo and Luoto, 2007). These signals are not easily discerned from static species distribution models and this hinders our ability to predict future species distributions. Dynamic models can allow us to detect how species interact with each other and how abiotic conditions mediate these interactions (Akesson et al. 2014). Our inverse modelling approach, combining *sed*aDNA plant community data with process-based models, could detect these signals, and supported the inclusion of temperature-mediated growth, competition between taxa through shading and top-down effects through the inclusion of an estimate of local herbivore pressure from reindeer. While the methods used would need to be adapted to reasonably scale up in terms of the number of taxa considered and for forecasting efforts, we find that they can give an ecologically relevant portrait that can complement statistical modelling efforts.

## Conflict of interest statement

The authors have no conflicts of interest to declare.

## Acknowledgements

We thank Dilli P. Rijal for sharing the interpolated summer temperature values for the sample points. Analyses were performed on computing resources provided by UNINETT Sigma2–the National Infrastructure for High-Performance Computing and Data Storage in Norway. The study was financed by The European Research Council (ERC) under the European Union’s Horizon 2020 research and innovation programme grant agreement No 819192 for the IceAGenT project (to I.G. Alsos) and The Arctic University Museum of Norway (supported M. Beaulieu).

## Author contributions

M.B., V.B., I.G.A. and L.P conceptualized the study. M.B., V.B. and L.P. developed the methodology. M.B. carried out the formal modelling analysis with input from V.B and L.P. M.B. interpreted the results and wrote the first version of the manuscript. V.B. L.P. and I.G.A. reviewed and edited the manuscript. All authors reviewed and approved the final version of the manuscript. I.G.A. acquired the funding that supported the project.

## Data Accessibility and Benefit-Sharing

Original plant data was published in Rijal et al. 2021 with re-analysing using the PhyloNorway DNA reference library (Alsos et al. 2022). The reindeer data were published in Alsos et al. (submitted). Julia code for inverse modelling and R code for data analysis can be accessed through github https://github.com/marieke-beaulieu/Dynamic-sedaDNA-models—Inverse-modelling

## Ethics Statement

This article reuses published data. Therefore, ethical approval was not required.

## Notes

### Competing Interest Statement

The authors have declared no competing interest.

### Summary of Updates

A better suited model performance metric was used to test training and testing performance.

