## Supplemental Figures for "Exploring the roles of temperature and competition in Fennoscandic Holocene plant communities using dynamic sedimentary ancient DNA models"

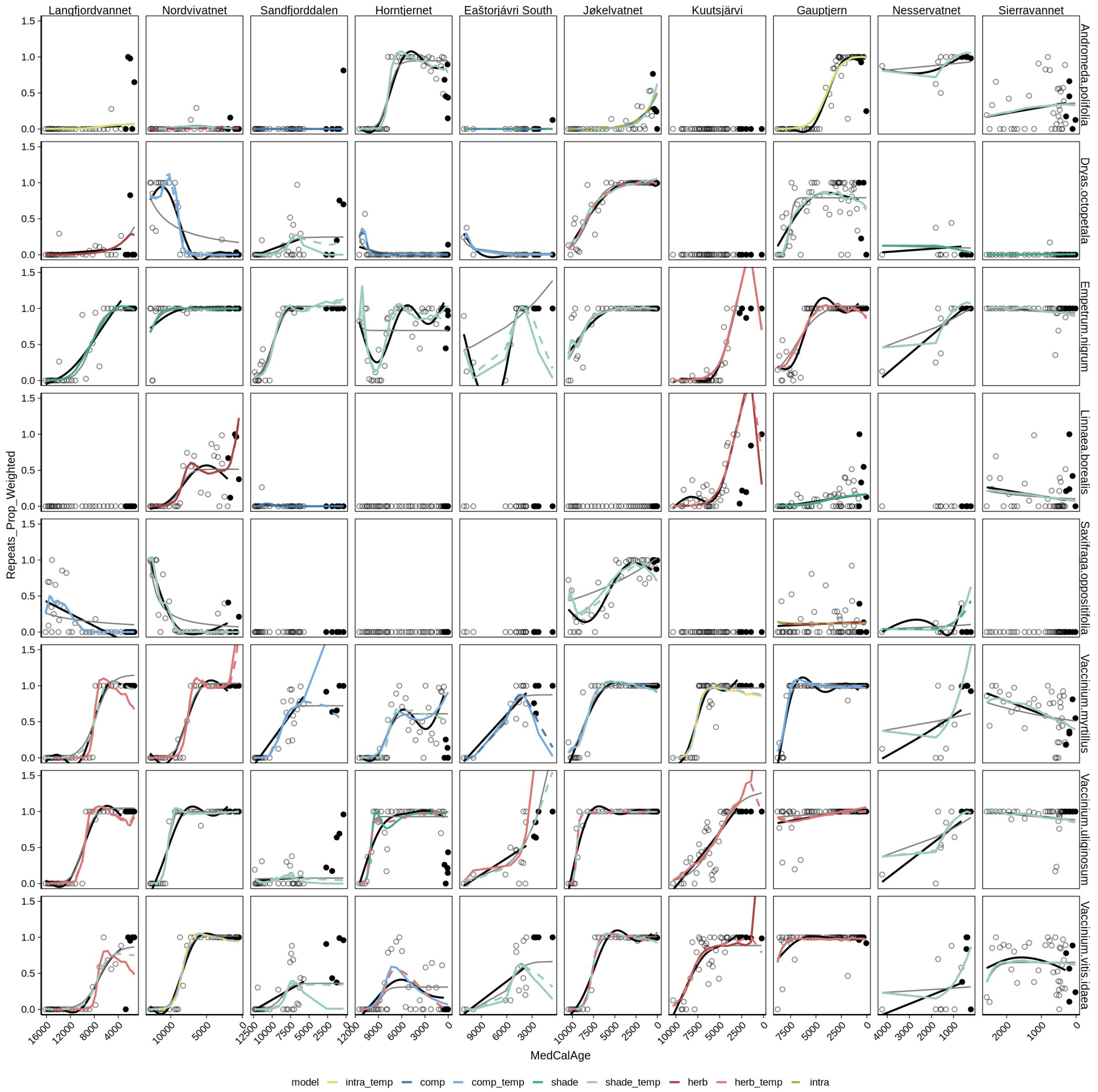


S1 Fitted model fits across selected lakes (columns) and taxa (rows). Baseline GAM model (k=5) shown in black. Null model (Basic intraspecific model) fit shown in grey. Best model based on training data MSE shown as coloured full line and best model based on testing data MSE from the top decile of trained models shown as broken coloured line.S2:

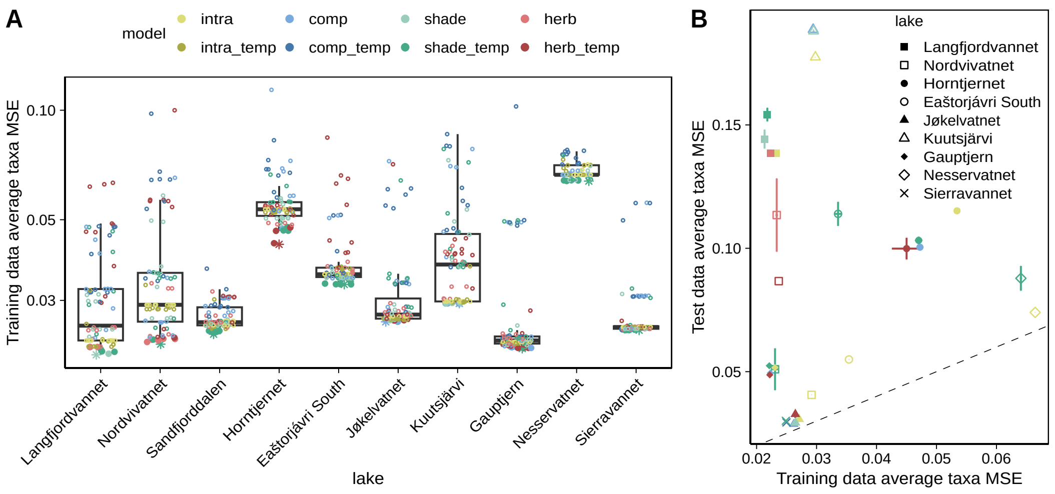


S2 Model fit. A: Community-wide training data fit as represented by the average taxon mean square error (MSE) for each model run (8 competing models over 5 runs, intra = intraspecific competition, comp = All taxa competition, shade= Shading competition, herb= Reindeer herbivory). Large filled points represent the top decile of models for each lake based on training MSE, with the star representing the best overall model. Open circles represent the remaining fitted models. B: Performance of the top decile of average taxa models based on the average taxa MSE of the testing dataset for each lake. For each lake, represented by a unique symbol, model runs were grouped for each of the model structures, using the same colours as in A) with crossbars showing the variance within. The null model (intra model1) is represented in yellow for comparison only and was never retained in the top decile of models. The dotted line represents the 1:1 ratio.

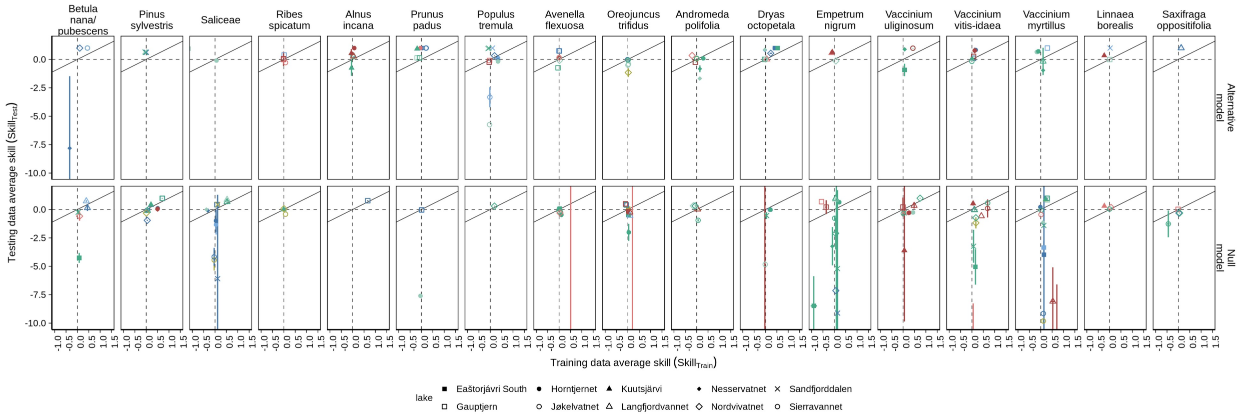


S3: Performance of the top decile of models for each taxa on the testing dataset. Top decile of models based on the Skill of the testing dataset for each lake. Data split on whether the model performed better on the test data than the null model (Alternative model) or if the null model fit the test data better (Null Model). When the null model was in the top decile of models it was included in the Null model group. Lakes represented as symbols and models represented as colours.
